# Temporal and spatial analysis of astrocytes following stroke identifies novel drivers of reactivity

**DOI:** 10.1101/2023.11.12.566710

**Authors:** Rachel D Kim, Anne E Marchildon, Paul W Frazel, Philip Hasel, Amy X Guo, Shane A Liddelow

## Abstract

Astrocytes undergo robust gene expression changes in response to a variety of perturbations, including ischemic injury. How these transitions are affected by time, and how heterogeneous and spatially distinct various reactive astrocyte populations are, remain unclear. To address these questions, we performed spatial transcriptomics as well as single nucleus RNAseq of ∼138,000 mouse forebrain astrocytes at 1, 3, and 14 days after ischemic injury. We observed a widespread and temporally diverse response across many astrocyte subtypes. We identified astrocyte clusters unique in injury, including a transiently proliferative substate that may be BRCA1-dependent. We also found an interferon-responsive population that rapidly expands to the perilesion cortex at 1 day and persists up to 14 days post stroke. These lowly abundant, spatially restricted populations are likely functionally important in post-injury stabilization and resolution. These datasets offer valuable insights into injury-induced reactive astrocyte heterogeneity and can be used to guide functional interrogation of biologically meaningful reactive astrocyte substates to understand their pro- and anti-reparative functions following acute injuries such as stroke.

## INTRODUCTION

Stroke is associated with neuronal death, vascular damage, and infiltration of peripheral immune cells – all of which provoke central nervous system (CNS) inflammation. Cell death is tightly regulated by inflammation following the initial event as well as in the subacute and chronic phases, with many pathological hallmarks such as glial scarring persisting for years^1^. Currently, fewer than 10% of patients receive effective therapeutics, namely thrombolytics, which are only clinically indicated to be administered within 4.5 hours post stroke^2^. Past research and therapeutic interventions for stroke have targeted neuronal vulnerability and death, but this approach has largely failed to reduce stroke-induced damage or improve functional recovery^3^. Looking beyond a neurocentric view to understand and target nonneuronal responses to injury is now crucial not only to understand the injury landscape at the molecular and functional levels, but also to identify novel therapeutic targets and treatments.

One such nonneuronal cell type of interest is astrocytes, which perform crucial developmental and homeostatic functions in the CNS^4^. Astrocytes, the most abundant nonneuronal cell type in the CNS, tile the entire brain and spinal cord and interface with a variety of neural and peripheral cells. These cells have rapid and robust responses to many insults, which result in transcriptomic and functional changes under a process broadly termed “reactivity”^5^.

Astrocyte reactivity is heterogeneous and depends contextually on the nature of the initiating insult^6^; notably, following acute CNS injuries such as stroke or spinal cord injury (SCI), a subpopulation of reactive astrocytes abutting the lesion core become transiently proliferative and give rise to the astrocyte scar border^7,8^. This border contains a chronic reactive astrocyte subtype that corrals proinflammatory peripheral immune cells to the necrotic lesion core^8^ and is typically not observed in other models of inflammation or neurodegenerative diseases. Because of the intrinsic heterogeneity in astrocyte responses to different insults, it is crucial to be able to identify insult-specific transient reactive astrocyte substates and chronic subtypes at the molecular, cellular, and functional levels.

Seminal *in vivo* microarray and bulk RNAseq experiments laid the groundwork for studying diversity in astrocyte reactivity at the gene expression level across different insults (including in acute injury models such as ischemic stroke^9-11^ and SCI^12^) but were not able to resolve diversity in astrocyte reactivity within the same insult. For example, previous *in situ* hybridization experiments showed that reactive astrocyte transcripts such as *Lcn2* and *Serpina3n* were upregulated only in astrocytes bordering the lesion in a mouse model of medial cerebral artery occlusion, contrasting with pan-tissue astrocyte upregulation of the same transcripts following broad inflammatory challenge with lipopolysaccharide (LPS) injection^11^. These findings demonstrated within-insult heterogeneity of reactive astrocytes – particularly following acute ischemic insult – and established the need for identification of multiple reactive astrocyte populations.

Single cell molecular profiling methods have now enabled further dissection of astrocyte substates^13-20^ within the same insult. However, many of these datasets use entire neural tissue as input rather than performing astrocyte enrichment. These datasets are valuable for resolving a broad image of the post-injury landscape, but for a still-unknown technical reason many single-cell technologies suffer from poor astrocyte capture, leading to the risk of underpowered astrocyte-specific data. This is particularly important as potentially rare but biologically meaningful reactive astrocyte substates may be masked. For example, our group previously enriched for – and sequenced – ∼80,000 astrocytes from LPS-injected mice with an average of 6,662 astrocytes per mouse and discovered a population of interferon-responsive reactive astrocytes (IRRAs) that comprised only 2.7% of all sequenced astrocytes^21^. Additionally, even with astrocyte-enriched single cell data, cell numbers captured can be too low to resolve sparse astrocyte clusters. This work demonstrates a need for astrocyte-specific enrichment for data that are sufficiently powered to identify lowly abundant reactive astrocyte substates at multiple timepoints following acute injury.

To better understand the molecular and temporal diversity of various astrocyte populations in response to injury, we set out to generate an astrocyte-enriched single transcriptomics dataset. We performed single nucleus RNAseq of ∼138,000 forebrain astrocyte nuclei isolated using fluorescence-activated nuclei sorting (FANS) from *Aldh1l1*-EGFP/ Rpl10a astrocyte reporter mice at 1, 3, and 14 days post stroke induced by Rose Bengal photothrombosis. To also resolve spatial heterogeneity of genome-wide gene expression in injured tissue, we performed spatial transcriptomics of lesioned brain slices at 1, 3, and 14d post stroke. Together, these datasets provide a powerful resource for probing injury-induced reactive astrocyte heterogeneity and can be used to guide functional interrogation of biologically meaningful reactive astrocyte substates to understand their potential pro- and anti-reparative functions following CNS injury.

## RESULTS

### Mouse forebrain astrocytes are molecularly and regionally distinct

To investigate molecularly and spatially distinct astrocyte subtypes, we used the Rose Bengal photothrombosis model of ischemic injury in 2-4 month old female and male *Aldh1l1*-EGFP/Rpl10a mice. At 1, 3, and 14d post stroke, we fresh froze ipsilateral forebrains for FANS isolation of GFP^+^ astrocyte nuclei (Figures 1A and S1A). Following quality control (see Methods), we were left with 151,917 total nuclei in 29 samples, 91% of which were astrocytes (Figures S1–S3), totaling 138,324 astrocytes and averaging 4,770 astrocytes per sample.

**Figure 1.**
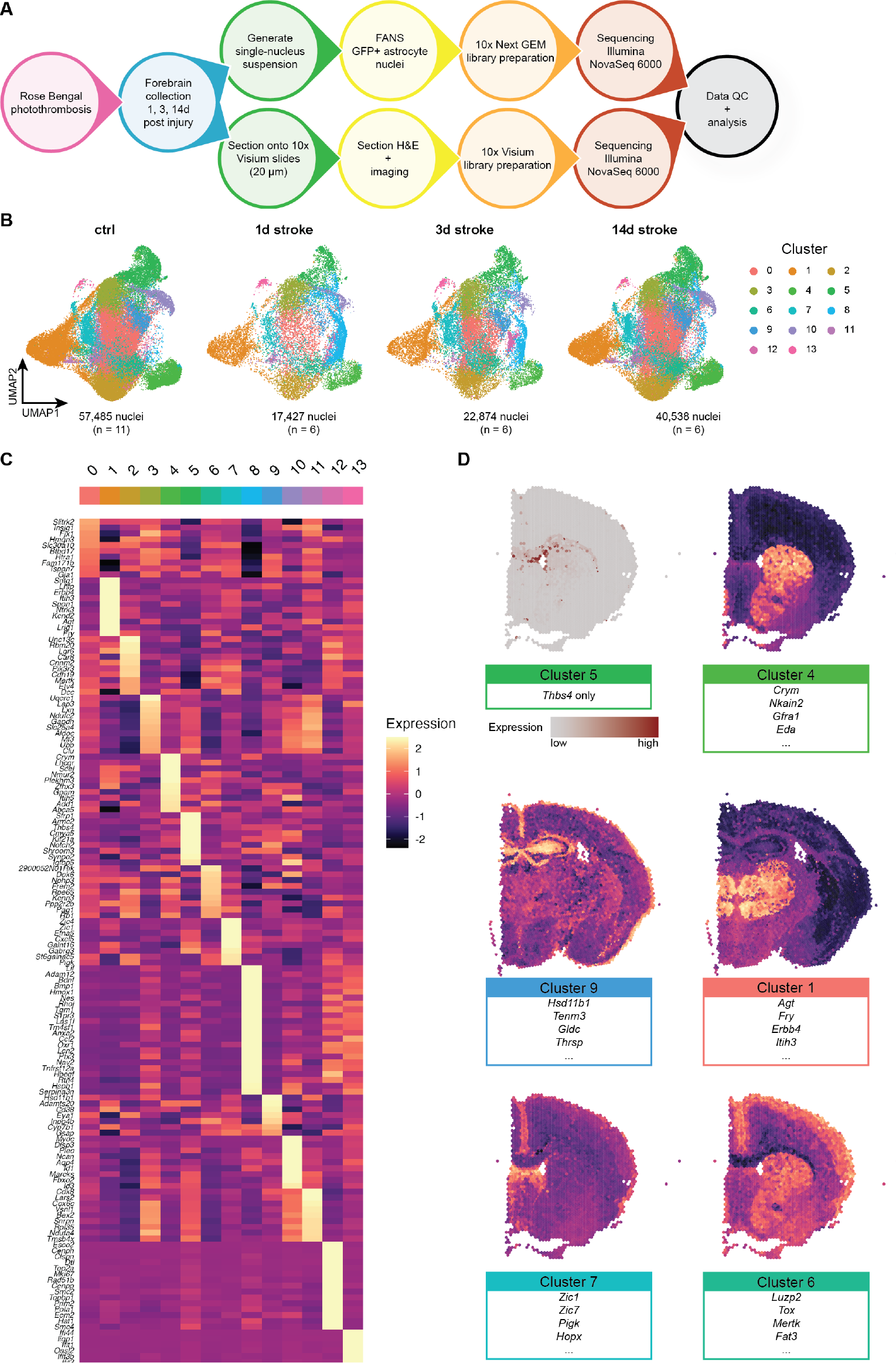
A large-scale astrocyte enriched single nucleus RNAseq dataset of molecularly and regionally distinct astrocyte subtypes. **(A)** Schematic of experimental approach for astrocyte-enriched snRNAseq and spatial transcriptomics of uninjured and injured brains using *Aldh1l1-*EGFP/Rpl10a mice. **(B)** After removal of non-astrocyte nuclei, 138,324 astrocytes were reclustered into 14 total clusters. UMAP plots are split across experimental condition: control (n = 5 male and 6 female mice), 1d post stroke (n = 3 male and female mice each), 3d post stroke (n = 3 male and female mice each), and 14d post stroke (n = 4 male and 2 female mice), from left to right. Individual astrocytes colored by cluster. **(C)** Heatmap of genes enriched in each astrocyte cluster. **(D)** Visium spatial transcriptomics of control tissue highlighting regionally distinct astrocyte clusters. Cluster 5 represents white matter astrocytes, which can be described by a single gene (*Thbs4*). We were able to infer localization of other astrocyte subtypes by using Seurat’s A*ddModuleScore* function.

We identified 14 molecularly distinct astrocyte clusters, many of which are enriched in previously identified (as well as new) marker genes (Figures 1B and 1C). For example, Cluster 1-enriched genes, including *Agt, Lhfp*, and *Itih3*, have been shown to be expressed in thalamic astrocytes^22^; Cluster 4 shows enrichment of striatal markers *Crym* and *Scel*^*2*3^; and Cluster 10, previously identified by our group as *glia limitans superficialis* astrocytes, which wrap the brain and spinal cord^24^, shows highly specific expression of *Myoc, Gfap*, and *Fbxo2*. We also identified clusters almost exclusively present in the injured brain. Cluster 8 is strongly enriched in broad reactive marker genes *Lcn2, Serpina3n, S1pr3*, and *Ptx3*^11^, as well as putative supportive factor genes *Lif, Bdnf*, and *Hbegf*; Cluster 12 is highly specific for cell cycle related genes such as *Mki67* and *Top2a*; Cluster 13 is enriched in interferon response genes, including *Oasl2, Gbp2, Gbp5, Gbp6*, and *Gbp7*.

To determine whether our astrocyte subtypes were regionally distinct, we performed genome-wide spatial transcriptomics on brain sections from two animals at each injury timepoint along with two corresponding control animals using 10x Genomics’ Visium platform (Figure S4). This method allows us to probe gene expression with spatial fidelity, and for astrocyte clusters whose marker genes are highly cell type- and astrocyte subtype-specific in an unbiased way. We were able to visualize where these astrocyte subtypes reside in the brain; notably, this can be observed in the *Thbs4*-expressing Cluster 5 white matter astrocytes^21^ (Figure 1D). Additionally, while this technology does not provide single-cell resolution, we were able to visualize groups of astrocyte cluster-enriched genes by analyzing their co-expression using Seurat’s *AddModuleScore* function (Figure 1D). For example, the Cluster 1 module (*Agt, Fry, Itih3*, etc.) shows enrichment in the thalamus, while the Cluster 4 module (*Crym, Scel, Eda*, etc.) is shown to be expressed most highly in the striatum, consistent with previous reports^23^. We also inferred the locations of Cluster 9 astrocytes (*Hsd11b1, Gldc, Thrsp*, etc.) to the hippocampus, Cluster 6 astrocytes (*Luzp2, Tox, Mertk*, etc.) to the upper cortex, and Cluster 7 astrocytes (*Zic1, Zic7, Hopx, Pigk*, etc.) to the lateral septum.

### Shared and subtype-specific reactive transformations following injury

To understand shared- and subtype-specific reactive transformations over time, we performed differential expression testing using muscat’s pseudobulk approach^25^ and observed robust gene expression changes across all clusters (Figures 2A and S5).

**Figure 2.**
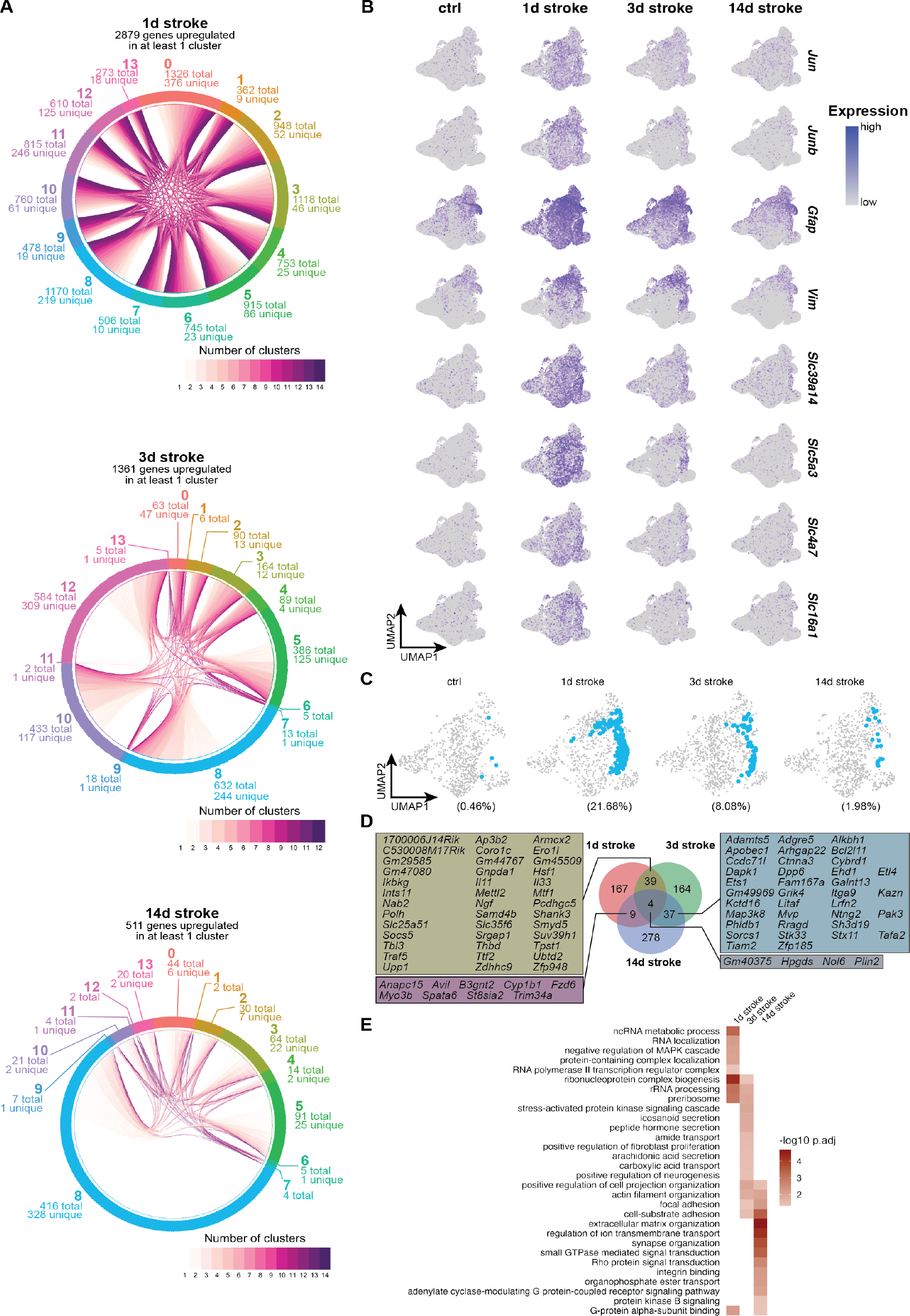
Shared and subtype-specific reactive transformations following injury. **(A)** Chord plot depicting shared and cluster-specific DEGs ≥ 1 log_2_ fold change and padj ≤ 0.05 at each timepoint. **(B)** Feature plots showing gene expression levels of individual genes identified as DEGs across all clusters at 1d post stroke, split by experimental condition. **(C)** UMAP plots highlighting cluster 8 (in blue), split by experimental condition. **(D)** Plot showing shared and overlapping DEGs unique to cluster 8 at each timepoint, and individual genes that are upregulated at multiple timepoints in cluster 8 astrocytes. **(E)** Heatmap showing GO terms associated with timepoint-specific upregulated DEGs in cluster 8 astrocytes.

Dramatic induction of differentially expressed genes (DEGs) was most notable at 1d post stroke, with hundreds of upregulated DEGs in all clusters (Figure 2A). Of the 2,879 genes upregulated at 1d post stroke, 71 genes were upregulated in all clusters. Among these 71 shared DEGs were early response genes *Jun* and *Junb*, which correlate with the acute phase of the injury (and have been reported previously^26-28^), as well as cytoskeletal components *Gfap* and *Vim*, which play roles in both extracellular^29^ and intracellular^30^ cell integrity. We also observed upregulation of many genes encoding solute carrier proteins; divalent metal transporter *Slc39a14*, myo-inositol transporter *Slc5a3*, sodium bicarbonate transporter *Slc4a7*, and monocarboxylate transporter *Slc16a1* (Figures 2B and S5C). The DEGs observed at 1d post stroke indicate that while there are many subtype-specific reactive responses, there is also a rapid shared reactive transformation to a state that may prioritize general maintenance of the environment after injury.

In most clusters at 3 and 14d post stroke there was a decrease in upregulated DEGs, suggesting a gradual return to transcriptional profiles that more resemble those of uninjured astrocytes (Figures 2A and S5C). There are a few exceptions to this pattern, including cluster 8 astrocytes, which represented a very sparse population of homeostatic astrocytes (0.46% of all uninjured astrocytes) that rapidly expanded around the lesion core at 1d (21.68% of all 1d stroke astrocytes) and gradually decreased up to 14d post stroke (8.08% and 1.98% of 3 and 14d stroke astrocytes, respectively; Figures 2C and S6) while maintaining hundreds of unique DEGs at their respective timepoints, even at 3 and 14d post stroke (Figures 2A, 2C—2E, S5C, S6, and S7).

At the early stages following injury (1 and 3d post stroke), cluster 8 astrocytes share upregulated DEGs coding for secreted signaling molecules such as *Il11, Il33*, and *Ngf*, as well as immune modulator *Socs5* and synaptic structure-related genes *Pcdhgc5* and *Shank3*. Cluster 8 astrocytes at these timepoints also upregulate stress response gene *Hsf1* as well as genes related to chromatin structure and regulation of cell cycle and mitotic transcription (*Polh, Samd4b, Smyd5, Suv39h1*, and *Ttf2*; Figures 2D and S7). Together, these data indicate multifaceted roles of cluster 8 astrocytes in interfacing with other cell types in the injured brain while tightly regulating their own internal transcriptional activity.

At later injury stages (3 and 14d post stroke), cluster 8 astrocytes share upregulated DEGs implicated with neurotrophic support, neurite outgrowth, and synapse formation (*Galnt13, Ntng2, Pak3, Tafa2*, and *Tiam2*). Additionally, we observed genes related to cell structure and extracellular matrix organization (*Adamts5, Ctnna3, Ehd1, Kazn, Kctd16, Phldb1, Sh3d19*, and *Stk33*; Figures 2D and S7). Upregulation of such genes may signal a functional transition to one that contributes to remodeling of injured tissue while remaining neurosupportive.

Cluster 8 astrocytes are not only distinct from other astrocyte clusters at their corresponding timepoints, but they are also transcriptionally different across time (Figure 2D). We performed functional gene annotation of timepoint-specific cluster 8 DEGs using clusterProfiler^31^ to broadly infer potential time-dependent functional transitions of cluster 8 astrocytes. At 1d post stroke, there is an enrichment of terms related to transcription and ribosome biogenesis; at 3d post stroke, there is an enrichment of terms related to cell morphology, interaction with other cell types, and transport and secretion of various substrates; at 14d post stroke there is an enrichment of terms regulated to physical cell-cell interactions and organization of extracellular space (Figure 2E). These results suggest that compared to other astrocyte clusters that undergo dramatic but transient transcriptional changes in response to injury, cluster 8 astrocytes sustain time-dependent substate transitions that may allow them to influence post-injury environment and repair.

### Cluster 12 – a spatiotemporally restricted substate associated with proliferation putatively regulated by BRCA1

In addition to the broad number of genes induced across multiple astrocyte clusters, we also observed lowly abundant but particularly interesting astrocyte clusters whose presence and gene expression profiles seem to be driven by injury. One such cluster is cluster 12, which expresses genes related to proliferation and cell cycle such as *Mki67* and *Top2a* (Figures 1C, 3A, and 3B). This cluster is specifically abundant at 3d post stroke (n = 513 total nuclei, 2.24% of all 3d stroke reactive astrocytes, and less than 1% of all astrocytes) with a gene expression signature present exclusively in this cluster (Figures 3A and 3B). Mapping the cluster 12 module onto Visium spatial transcriptomic data shows expression of this module in regions abutting the lesion core only at 3d post stroke (Figure 3C). Validation at higher resolution using *in situ* hybridization indicates colocalization of the proliferation marker *Top2a* and astrocyte marker *Slc1a3* in astrocytes abutting the lesion core only at 3d stroke (0.122 ± 0.109, 4.332 ± 0.872, and 0.663 ± 0.388% of *Slc1a3*^+^ astrocytes per hemibrain at 1, 3, and 14d post stroke, respectively; Figures 3D, 3E, and S8). These data confirm the presence of transiently proliferative astrocytes following Rose Bengal photothrombosis and are consistent with previous reports in other injuries such as spinal cord injury (SCI) and stab injury, which report proliferation at 3-5 days post injury of astrocytes neighboring the lesion core^7,8^.

**Figure 3.**
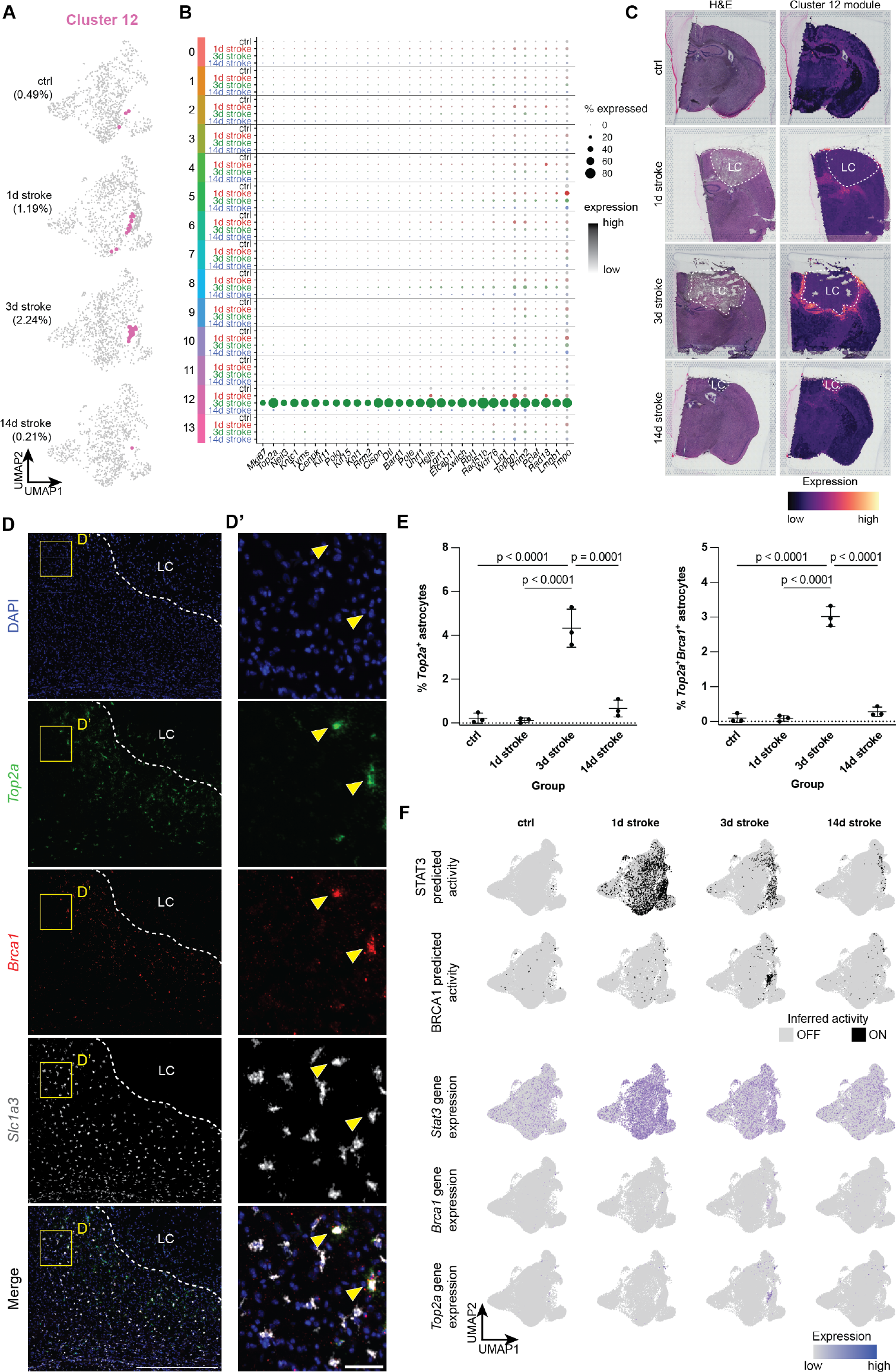
Cluster 12 – a tightly spatiotemporally restricted substate associated with proliferation and may be regulated by *Brca1*. **(A)** UMAP plots highlighting cluster 12 (in pink), split by experimental condition. **(B)** Dot plot showing cluster 12-enriched genes, split across all clusters by experimental condition (gray: control, red: 1d post stroke, green: 3d post stroke, blue: 14d post stroke). **(C)** Visium spatial transcriptomics highlighting cluster 12-enriched genes across all experimental conditions. Left: H&E stain of individual sections, Right: cluster 12 gene module expression. LC = lesion core. **(D)** Representative image of 3d post stroke section for nuclei (DAPI), proliferation marker *Top2a, Brca1*, and astrocyte marker *Slc1a3*. Scale = 500 μm **(D’)** Zoomed image from Panel E. Scale = 50 μm. **(E)** Quantification of % *Top2a*^+^ astrocytes as well as *Top2a*^+^*Brca1*^+^ astrocytes. Data shown as mean ± SD. Statistical analyses were performed by a one way ANOVA with Bonferroni’s multiple comparison. **(F)** Above: plots showing inferred on/off (on: black, off: grey) states of transcriptional regulators *Stat3* and *Brca1*. Below: Feature plots showing expression of *Brca1* and proliferation marker *Top2a*, showing expression only in cluster 12 astrocytes at 3d post stroke.

We next sought to infer transcriptional regulators for cluster 12 proliferation-associated astrocytes using gene regulatory network analysis (GRN) with pySCENIC^32^. Foundational work has shown that this population of reactive astrocytes was reliant on the transcription factor STAT3, and that conditional knockout of *Stat3* from *Gfap*^+^ astrocytes disturbed formation of the astrocyte scar border and allowed for increased infiltration of proinflammatory peripheral immune cells into the parenchyma^8,33^. Our analysis inferred the activation of STAT3 not only in cluster 12 astrocytes, but also across all astrocyte clusters at 1d post stroke. This predicted activation becomes restricted to fewer clusters at 3 and 14d post stroke but is not necessarily cluster 12-specific, which is also reflected at the gene expression level (Figures 3F and S9A). This suggests that STAT3 is a potent but promiscuous transcriptional regulator especially at 1d post stroke, raising the concern that perturbation of STAT3 signaling may lead to yet undescribed influences on reactive astrocyte substates outside of the astrocyte scar border. Instead, we identified several transcriptional regulators, the gene expression and inferred activation of which are enriched in 3d stroke cluster 12 astrocytes, including BRCA1, E2F1/2/7/8, and MYBLl1/2 (Figure S9B). We found in our snRNAseq data that of these predicted transcriptional regulators, *Brca1* was highly expressed almost exclusively in 3d stroke cluster 12 astrocytes (Figure 3F). *In situ* hybridization of *Brca1, Top2a*, and *Slc1a3* confirms that *Top2a* and *Brca1* are indeed co-expressed in *Slc1a3*^+^ astrocytes abutting the lesion core only at 3d post stroke (0.091 ± 0.085, 3.018 ± 0.286, and 0.283 ± 0.136% of *Slc1a3*^+^ astrocytes per hemibrain at 1, 3, and 14d post stroke, respectively; Figures 3D, 3E, and S8). We therefore suggest that *Brca1* may be a more subtype- and substate-specific transcriptional regulator of proliferative reactive astrocytes, which have been shown to give rise to the astrocyte scar border following other models of CNS injury^7,8^.

### Cluster 13 – interferon-responsive reactive astrocytes (IRRAs) rapidly expand and persist cortically following stroke

Another prominent reactive astrocyte cluster of particular interest was cluster 13, which our group has previously characterized as interferon-responsive reactive astrocytes^21^ (IRRAs; Figures 4A-4C) that are closely associated with blood vessels, ventricles, and the brain surface. Unlike temporally transient proliferative cluster 12 astrocytes, IRRAs reside in the white matter and ventricles in the healthy brain (0.26% of all control astrocytes; Figures 4D and 4E). Following injury, IRRAs rapidly expand to occupy the perilesion at 1d post stroke (1.61% of all 1d stroke astrocytes) and persist up to 14 days (0.56% of all 14d stroke astrocytes; Figure 4F). Mapping the cluster 13 module on Visium data shows expression of this module in spots around the lesion core only at 1 and 3d post stroke. Interestingly, in 14d stroke Visium tissue, cluster 13 module expression is highest inside the lesion core (Figure 4D). However, *in situ* hybridization for IRRA marker Igtp and astrocyte marker *Slc1a3* shows colocalization of *Igtp* and *Slc1a3* at all timepoints around the lesion core. These data confirm not only our snRNAseq findings of white matter- and ventricle-associated IRRAs in the healthy brain (0.76 ± 0.16% of *Slc1a3*^+^ astrocytes per hemibrain), but also the expansion of IRRAs occupying areas surrounding the lesion (2.74 ± 0.60, 2.40 ± 0.44, and 1.14 ± 0.36% of *Slc1a3*^+^ astrocytes per hemibrain at 1, 3, and 14d post stroke, respectively; Figures 4E and 4F).

**Figure 4.**
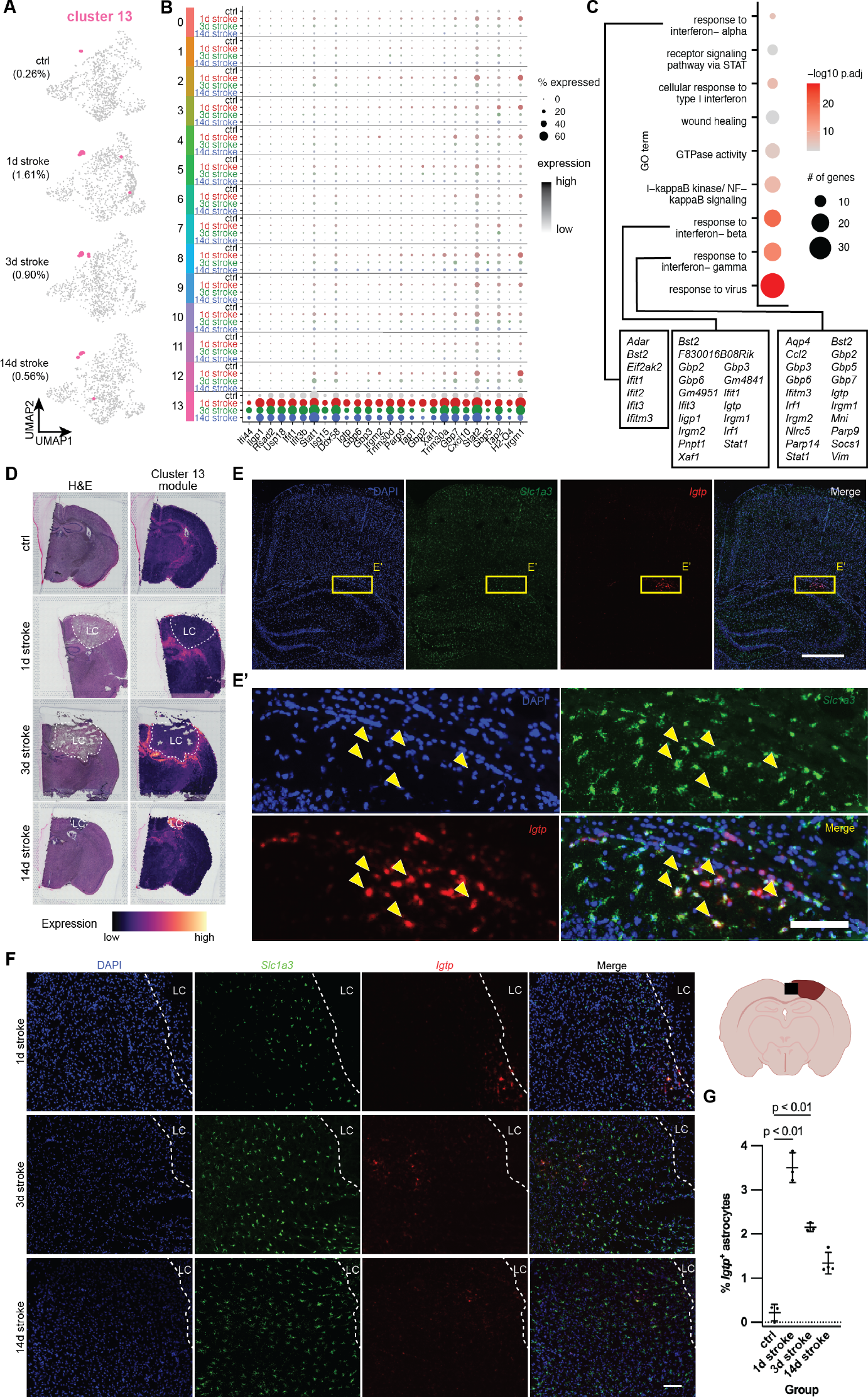
Cluster 13 – interferon-responsive reactive astrocytes (IRRAs) are rapidly expanded in perilesion cortex following stroke and persist chronically. **(A)** UMAP plots highlighting cluster 13 (in pink), split by experimental condition. **(B)** Dot plot showing cluster 13-enriched genes, split across all clusters by experimental condition (gray: control, red: 1d post stroke, green: 3d post stroke, blue: 14d post stroke). **(C)** Dot plot showing GO terms of cluster 13-enriched genes. Dot size indicates number of genes, color represents padj. In text boxes are cluster 13-enriched genes in GO terms related to interferon signaling. **(D)** Visium spatial transcriptomics highlighting cluster 13-enriched genes across all experimental conditions. Left: H&E stain of individual sections, Right: cluster 13 gene module expression. LC = lesion core. **(E)** Representative in situ image of control section for nuclei (DAPI), IRRA marker *Igtp*, and astrocyte marker *Slc1a3*. Scale = 500 μm. **(E’)** Zoomed image from Panel E. Scale = 100 μm. **(F)** Representative in situ images at 1, 3, and 14d post stroke cortex for nuclei (DAPI), IRRA marker *Igtp*, and astrocyte marker *Slc1a3*. Scale = 100 μm. **(G)** Quantification showing % of *Igtp*+ astrocytes in the ipsilateral hemispheres of 1, 3, and 14d post stroke tissue. Data shown as mean ± SD. Statistical analyses were performed by a one way ANOVA with Bonferroni’s multiple comparison.

## DISCUSSION

Descriptions of reactive astrocytes following injury have long been limited by the profiling methods used (microarray and bulk profiling vs. single cell profiling). This is particularly notable in a perturbation such as acute injury, where there are distinct compartments in the injury environment (a necrotic lesion core, plastic perilesion, spared tissue, etc.) that is highly dynamic over time. Additionally, many single-cell interrogations of CNS injuries use whole tissue as input and suffer from poor astrocyte capture; this leads to insufficient power required to bioinformatically identify rare but potentially biologically meaningful astrocyte subtypes and reactive substates. To address these limitations, we combined multimodal transcriptomic profiling (snRNAseq and spatial transcriptomics) and astrocyte enrichment using FANS, and we found that the reactive astrocyte landscape after ischemic injury is highly diverse and dynamic in various astrocyte subtypes.

Transcriptomically, most astrocytes cluster by brain region, consistent with previous studies probing homeostatic astrocyte heterogeneity with both bulk^23,34,35^ and single-cell^21,24,36^ methods. After injury, we uncovered complex and dynamic cluster- and time-dependent changes in gene expression across all astrocyte clusters, with 1d stroke astrocytes dramatically upregulating hundreds of DEGs across all clusters. Of all these genes, 71 were significantly upregulated in all 14 clusters; this included early response genes *Jun* and *Junb* and radial glial/astrocyte markers *Gfap* and *Vim*. Additionally, all astrocyte clusters upregulated various solute carrier transport genes at 1d stroke. Manganese transporter *Slc39a14*, whose knockout of which in mice leads to accumulation in manganese accompanied by upregulation of various proinflammatory transcripts and decreased motor activity^37^; *Slc5a3*, a myo-inositol transporter, which has been theorized to provide neuroprotection following injury^38^; sodium bicarbonate transporter *Slc4a7*, which has been shown *in vitro* to be essential for astrocyte survival in ischemic conditions^39^ and acts as a protective mechanism from acidification^40^; and *Slc16a1*, a monocarboxylate transporter that provides crucial metabolic support, especially to neurons^41,42^. Altogether, upregulation of these transporters seems to indicate a broad, injury compartment-independent response that is dedicated both to supporting other cells and altering the extracellular environment in the injured nervous system.

We also observed that the number of upregulated DEGs decreased across almost all clusters over time, suggesting that most reactive substates are transient and can gradually revert to a state that is more similar to an uninjured homeostatic state. A notable exception to this observation is cluster 8, which remains transcriptionally distinct from its uninjured state even at 14d post stroke. Gene expression within this cluster is highly dynamic and diverse over time. This suggests that the injury environment is highly plastic, and that there remains a population of astrocytes that mirrors this plasticity at the gene expression level. To what environmental signals and cell-cell interactions cluster 8 astrocytes are responding and affecting, as well as what cluster 8 astrocyte functions are, remains elusive. This now raises the question of if, and how, the injured nervous system is able to return to an uninjured state at timepoints that are more chronic than 14d post stroke.

We highlight two transcriptionally distinct substates. One (cluster 12) is a rare population of proliferative astrocytes that is tightly spatiotemporally restricted. We posit that this is likely the population of reactive astrocytes that gives rise to the scar border and whose organization is perturbed by *Stat3* knockout in *Gfap*^+^ astrocytes^8,12,33^. Our analysis suggests that while a powerful driver of this cluster, STAT3 may also drive transcriptional changes in many other astrocyte populations, potentially making it too broad a driver for investigating the function of specific substates of reactive astrocytes. We instead identified BRCA1 as a potentially more specific regulator of cluster 12 reactive astrocytes. We used *in situ* hybridization to confirm their gain of proliferative function, and demonstrated that *Brca1* is colocalized with *Top2a* only in 3d stroke reactive astrocytes in the peri-infarct region. It is unclear whether BRCA1 itself is a direct driver of cluster 12 astrocytes, and whether it interacts at all with STAT3. Further, the implications of BRCA1 conditional ablation from astrocytes for scar border formation and functional recovery from stroke and other insults where a proliferative scar-border is putatively helpful for resolution of the insult is unknown.

Lastly, we describe cluster 13 IRRAs and their presence in the post-stroke perilesion. We previously identified IRRAs in rodent models of general inflammation and neurodegenerative disease^21^. Outside models of broad inflammation or neurodegenerative diseases, IRRAs have been observed in multiple injury datasets^9,16,43^. What remained uncertain was when this population is induced after injury, whether this is a stable population, and what signaling pathways and mechanisms control this population. In this study, we show that IRRAs are a lowly abundant, spatially restricted astrocyte population that expands to the cortical perilesion at early timepoints (1d post stroke) and persist up to 14 days. Interestingly, spatial transcriptomics of this interferon response gene signature show highest expression inside the necrotic lesion core at 14d post stroke. We posit that this interferon response signature can also be induced in many other cell types; for example, fibrotic scar-forming fibroblasts inside the lesion core depends on type II interferon signaling^44^. This vignette is a compelling reminder that our spatial transcriptomics – while a powerful tool for whole-genome exploration of spatial compartments – is not cell type specific, and that *in situ* validation of sequencing data is crucial. Using gene ontology, we also propose that this subtype is regulated by type I or type II interferon signaling. Whether perturbation of type I and II interferon signaling affects formation and function of IRRAs remains unknown, as well as what cell type or cell types could be responsible for producing these interferons. Flow cytometry data of immune cells following ischemic injury reveal various populations of immune cells infiltrating the injured brain^45,46^, including type I interferon-producing macrophages and type II interferon-producing Natural Killer T cells^47^. The degree to which IRRAs participate in events following injury remains uncertain: do they influence or interact with not only other cell types, but also with other reactive astrocyte subtypes or substates such as proliferative scar-forming reactive astrocytes?

Combined, these data provide comprehensive insight into the diversity and dynamism of injury-induced reactive astrocyte subtypes and substates at the single cell transcriptional level. We hope that that these data can be used not only for bioinformatic comparisons but also as a springboard for hypotheses and further functional interrogations of these reactive populations.

## METHODS

### Animals

All animal experiments were approved by NYU Grossman School of Medicine’s institutional Animal Care and Use Committee (IACUC). For snRNAseq, Aldh1l1-EGFP/Rpl10a (B6;FVB-Tg(Aldh1l1-EGFP/Rpl10a)JD130Htz/J, RRID: IMSR_JAX:030247) were used. For spatial transcriptomics and in situ imaging, WT C57BL/6J mice (RRID: IMSR_ JAX:000664) were used. Mice were housed on a 12 h light/ dark cycle and were given food and water *ad libitum*. All animal procedures were in accordance with the guidelines provided by the National Institute of Health as well as NYU School of Medicine’s Administrative Panel on Laboratory Animal Care. All animals were housed at 22–25 °C and 50–60% humidity.

### Rose Bengal photothrombosis

All procedures were performed under general anesthesia with isoflurane (3-4% isoflurane with 0.25-1 L/min oxygen in induction chamber, 1-3% isoflurane with 0.25-1 L/min oxygen using a nose cone for maintenance) and a rodent stereotaxic frame. 2-5 month female and male mice were weighed and injected i.p. with 15 mg/kg Rose Bengal (Sigma-Aldrich 330000) dissolved in physiological saline (Addipak 200-59), as well as an s.c. injection of 5 mg/kg Ketoprofen (Zoetis) for surgical analgesia. The skin above the cranium was clipped, disinfected with betadine, and a small midline incision was made using a scalpel to expose the skull. After 5 min to allow diffusion of Rose Bengal throughout the bloodstream, a cold light source (150 W, Schott KL 1600 LED) was placed over a stainless steel ring (hole diameter 3 mm) on top of the animal’s skull and light stimulation was applied for 10 min. After light exposure, tissue adhesive (WPI VETBOND) was applied to the edges of the incision. Animals were returned to a clean cage over a heat pad and monitored until fully recovered from anesthesia; for post-operative care, animals were monitored daily and given 5 mg/kg Ketoprofen as instructed.

### Tissue collection, nuclei isolation, and fluorescence activated nucleus sorting (FANS) for snRNAseq

Animals were euthanized by rapid decapitation at 1, 3, and 14d post stroke and brains were extracted and moved to cold 1x PBS (Thermo Fisher SH30256.FS). Olfactory bulbs, cerebellum, and midbrain were first removed, then the ipsilateral and contralateral hemispheres were separated and stored in -80 °C until nuclei isolation.

Nuclei isolation was adapted from a previously published isolation protocol^48^ and completed on ice or at 4 °C. All solutions were prepared in a base dissociation buffer (82 mM Na-_2_SO_4_, 30 mM K_2_SO_4_, 10 mM glucose, 10 mM HEPES, 5 mM MgCl_2_.6H_2_O in water). Ipsilateral hemispheres were minced using a scalpel and moved to a well of a pre-chilled 6 well plate containing an extraction buffer containing of 1% Kollidon VA64, 1% Triton X-100 (Sigma-Aldrich T8787), 1:100 RNasin Plus (Promega N2615), and 1:200 Alexa Fluor 488 anti-GFP antibody (BioLegend 338007) in dissociation buffer. Samples were then mechanically triturated using a P1000 pipette and incubated for 10 min with occasional mechanical trituration. Samples were then pulled into a chilled 5 mL syringe/27 G needle and expressed into a second, then third well. After filtering, the single nucleus suspension was spun for 10 min at 600 g in a pre-chilled centrifuge. The nuclear pellet was resuspended and filtered, then spiked with TOPRO3 (Thermo Fisher R37170) to visualize nuclei.

Nuclei sorting was performed on a Sony SH800Z with a 100 μm nozzle at 4 °C (see Figure S1A for representative gating scheme). TO-PRO3^+^GFP^+^ nuclei were collected in 1:40 RNasin Plus in dissociation buffer. After sorting, collected nuclei were pelleted at 600 g for 10 min at 4 °C, and resuspended in ∼100 μL of dissociation buffer.

### 10x snRNAseq pipeline

Nuclei were prepared using the 10x Chromium Next GEM Single Cell 3’ kit (v3.1) according to the manufacturer’s instructions. Nuclei were loaded onto the Chromium Next GEM chip with the goal to recover 5,000-10,000 nuclei per sample. Gel bead-in emulsions were created using the Chromium Controller, following by cDNA synthesis, amplification, and library construction following the 10x pipeline. Quality and concentration of cDNA were evaluated on an Agilent 2100 Bioanalyzer. Quality and concentration of libraries were evaluated by qPCR and on an Agilent 2200 TapeStation, and libraries were sequenced an Illumina NovaSeq 6000 through NYU Langone’s Genomic Technology Core. Using Cell Ranger software suite (v6.0.1) (10X Genomics), FASTQ files were aligned to a pre-mRNA-modified mm10-2020-A mouse reference genome, and gene-barcode count matrices were generated for all demultiplexed samples.

### snRNAseq analysis

#### QC and processing

CellBender^49^ (0.2.0) was used to correct for ambient/background RNA using the Cell Ranger raw .h5 file as input. Doublet detection and removal was performed using DoubletFinder^50^ (2.0.3), and outliers were removed using scuttle^51^ (1.4.0). Using Seurat^52^ (4.3.0), data were log normalized and the 2,000 most variable features were identified for each sample. Samples were anchored and batch corrected using canonical correlation analysis (dims = 30). The batch corrected object was scaled, and linear and nonlinear dimensional reduction was performed using principal component analysis (PCA) and uniform manifold approximation and projection (UMAP), respectively.

#### Clustering

Coarse cell type clusters were identified using Seurat’s *FindClusters* function at a resolution of 0.1 and were able to identify clear clusters that expressed various canonical marker genes. We subsequently subsetted astrocytes for downstream analysis and performed a second iteration of batch correction and clustering using the same pipeline described above.

#### Differential gene expression

For differential state testing of astrocyte clusters across experimental conditions, the muscat^25^ (1.8.2) pipeline was used. In brief, sample level data for each cluster were aggregated to generate pseudobulk data. Then edgeR^53^ (3.36.0) was used to calculate differentially expressed genes (DEGs) among experimental conditions on the sum of counts across all samples for each experimental condition for each cluster; the threshold for significance was log_2_FC ≥ 1 and padj. ≤ 0.05. clusterProfiler^31^ (4.2.2) was employed for gene functional annotation of statistically significant DEGs, using FDR ≤ 0.05 as cutoff.

#### Inference of transcriptional regulator activity

To infer transcriptional regulators of astrocytes, we used the pySCENIC^32^ (0.11.2). In brief, sets of genes that were coexpressed with known transcription factors were identified as coexpression modules. Then, cis-regulatory motif analysis was used to identify putative direct-binding targets and filter for significant motif enrichment. To infer “ON”/”OFF” states of transcriptional regulators for each astrocyte we then used pySCENIC’s *AUCell* algorithm.

### Spatial transcriptomics

#### Sample processing

For spatial transcriptomics we used the 10x Visium platform (v1) following the manufacturer’s instructions. In short, fresh frozen brains sections for control and 1, 3, and 14d stroke mice were coronally sectioned at 20 μm onto Visium capture slides (n = 2 for each experimental condition). Following methanol fixation at -20 °C for 30 min, sections were stained with hematoxylin and eosin, then imaged at 20x and tiled using a Keyence BZ-X710 and BZ-X710 software, respectively. Sections were then enzymatically permeabilized for 14 min to add spatial barcodes and unique molecular identifiers to the captured polyA mRNAs. Libraries were generated using Dual Index primers and sequenced using an Illumina NovaSeq6000. SpaceRanger (1.3.1) was then used to align the sequencing output to the mm10 mouse reference genome as well as for aligning sequenced spot data to section image data.

Further analysis was performed using Seurat (4.3.0) where each section was transformed using Seurat’s *SCTransform* function, then merged and clustered based on variable features from each sample.

#### Subspot resolution using BayesSpace

For enhancing resolution of 10x Visium data, we used the BayesSpace^54^ (1.4.1) pipeline. In brief, individual SCE objects were created from the merged Seurat object. Subspot resolution was enhanced using the *spatialEnhance* function, then spatial gene expression for all genes was enhanced to subplot resolution using the enhanceFeatures function. To highlight astrocyte clusters identified in our snRNAseq data, Seurat’s *AddModuleScore* function was used.

### Tissue collection and RNA in situ hybridization

#### Sample processing

Mice were sacrificed at either 1, 3, or 14d post stroke with their respective controls. Brains were dissected and placed in Tissue-Tek O.C.T (Sakura Finetek USA 4583) in cryomolds (Sigma-Aldrich E6032) and frozen in dry ice and isopropanol. Samples were placed in a Leica cryostat one hour prior to sectioning and sectioned at 20 μm onto Superfrost plus microslides (VWR, 48311-703). Samples were stored in -80 °C until use.

*In situ* hybridization was performed using the ACDBio RNAscope Multiplex Fluorescent Reagent Kit (v2) according to the manufacturer’s instructions. Slides were placed immediately into 4% paraformaldehyde in 4 °C for 15 min. They were rinsed twice with PBS and underwent dehydration with ethanol at 50%, 75%, and two 100% washes for 5 min each. Samples were allowed to dry at room temperature. Lipid barriers were drawn with an Immedge pen (ACD, 310018). Samples were incubated with hydrogen peroxide at room temperature for 15 min, followed by two washes with water, and Protease IV was added for 30 min at room temperature. The protease was washed twice with PBS, and the following probes were added: *Top2a* in C1, *Igtp* in C1, *Brca1* in C2, and *Slc1a3* in C3. The probes incubated for 2 h in 40 °C in the HybEz oven. Samples were washed with wash buffer two times for 2 min and then amplified according to the V2 protocol. Opal fluorophores 520 (Akoya Biosciences FP1488001KT), 620 (Akoya Biosciences FP1495001KT) and 690 (Akoya Biosciences FP1497001KT) were added at a 1:1000 dilution to their corresponding probes according to the protocol, and counterstained with DAPI. Fluoromount-G (SouthernBiotech, 0100-01) was used to mount the slides, and they were covered with a 50 mm cover glass slip (Denville Scientific M1100-02) and left at room temperature in the dark overnight to dry.

#### Quantification

Quantification of images acquired with in situ hybridization was done using QuPath^55^ (v0.4.3). Individual nuclei were detected based on DAPI signal. Duplicate channel training images were then generated to train an object classifier in each channel (*Slc1a3* and *Igtp* for cluster 13 images and *Slc1a3, Top2a*, and *Brca1* for cluster 12 images). Object classifiers were then loaded onto the merged images to generate cell counts.

## Acknowledgements

R.D.K. was supported by an NSF Predoctoral Fellowship. S.A.L. was supported by the NIH National Eye Institute (R01EY033353), NYU Grossman School of Medicine, the Neurodegenerative Diseases Consortium from MD Anderson, the Gifford Family Neuroimmune Consortium as part of the Cure Alzheimer’s Fund, the Carol and Gene Ludwig Family Foundation, the National Multiple Sclerosis Society, generous anonymous donors, and the financial support of Paul Slavick. The computational requirements for this work were supported in part by the NYU Langone High Performance Computing (HPC) Core’s resources and personnel as well as the NYU Langone Flow Cytometry Core. We also acknowledge the shared resources of the NYU Langone Genome Technology center that is partially supported by a Cancer Center Support Grant (P30CA016087) at the Laura and Isaac Perlmutter Cancer Center.

## Author Contributions

Conceptualization, R.D.K. and S.A.L.; Methodology, R.D.K. and S.A.L.; Software, R.D.K; Validation, R.D.K. and A.E.M; Formal Analysis, R.D.K.; Investigation, R.D.K., A.E.M., P.W.F., P.H., and A.X.G.; Data Curation, R.D.K.; Writing – Original Draft, R.D.K., A.E.M., S.A.L.; Funding Acquisition: R.D.K. and S.A.L.; Visualization, R.D.K; Supervision, S.A.L. All authors edited and approved the final draft of the manuscript.

---

**Figure S1.**
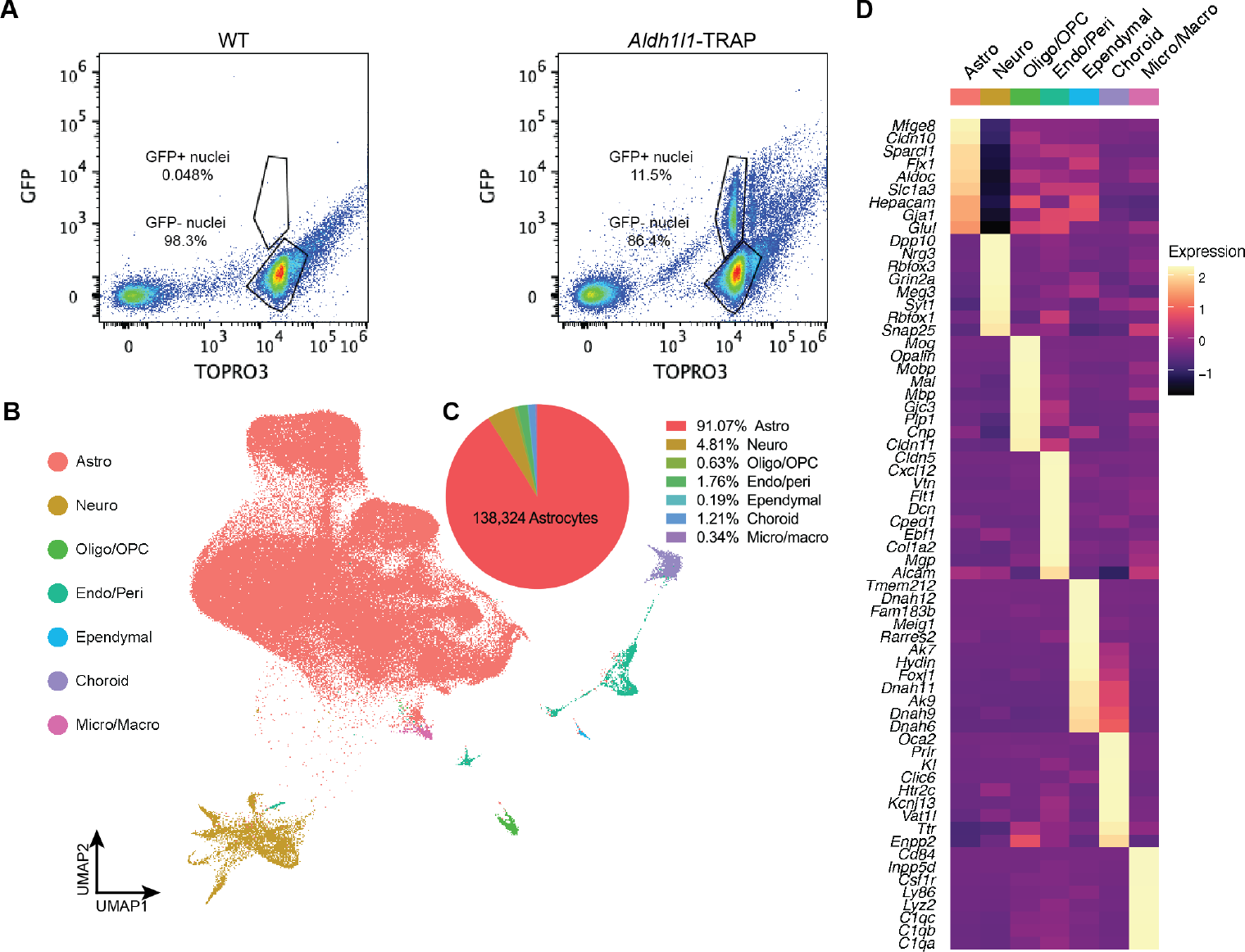
Astrocyte enrichment using *Aldh1l1*-EGFP/Rpl10a mice with FANS. **(A)** Sorting strategy for astrocyte enrichment with WT vs *Aldh1l1*-EGFP/Rpl10a mice. **(B)** UMAP plot of all sequenced nuclei across 29 samples. Following initial quality control, 151,891 total cells were retained, with astrocytes representing 91.07% of all high quality sequenced nuclei. **(C)** Pie chart depicting percentage of cell types identified in the dataset. **(D)** Heatmap of genes enriched in each cell type.

**Figure S2.**
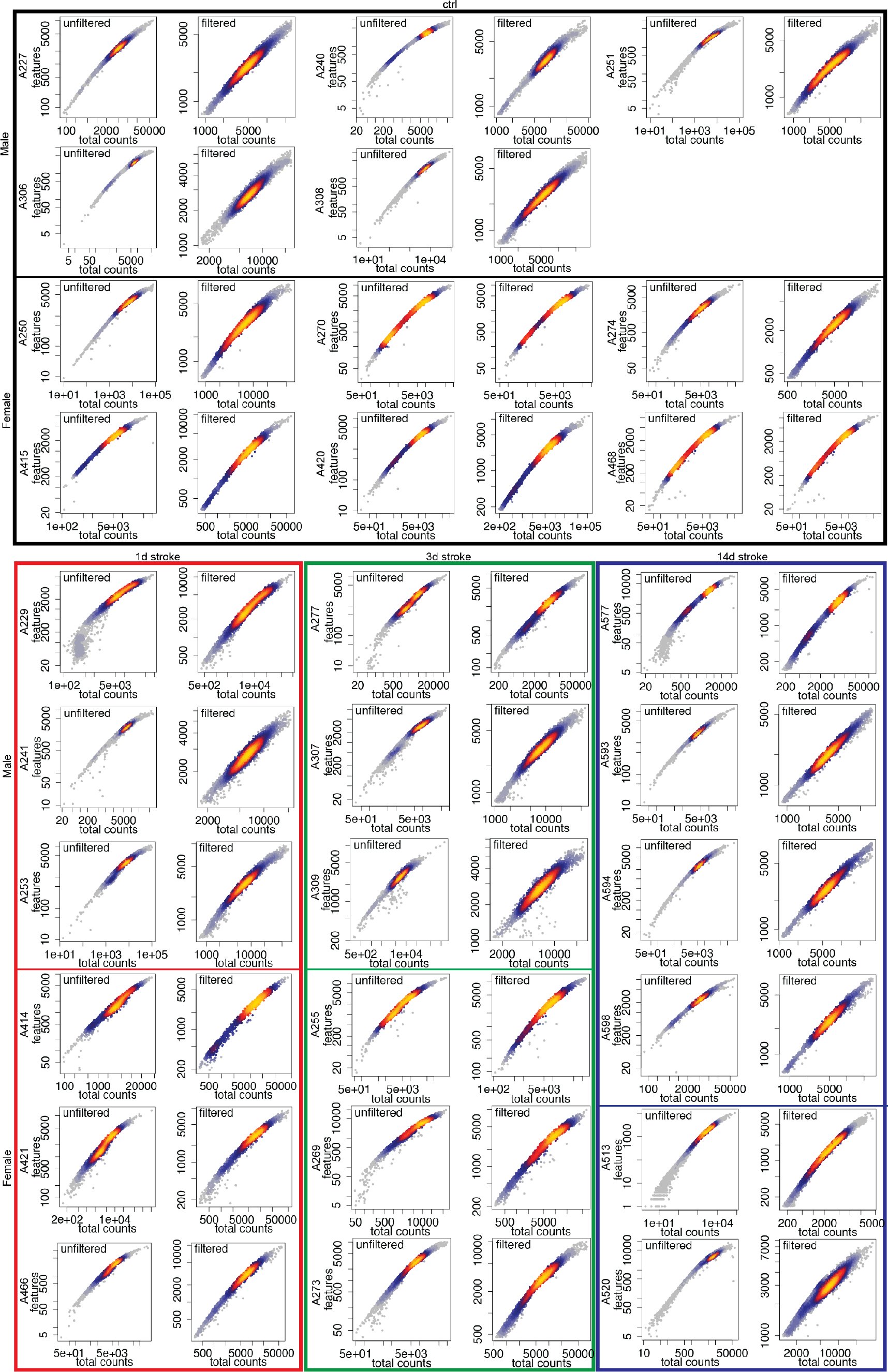
QC of nuclei collected for snRNAseq. Scatter plots showing counts (x-axis) and features (y-axis) for each sample, before and after QC/outlier removal.

**Figure S3.**
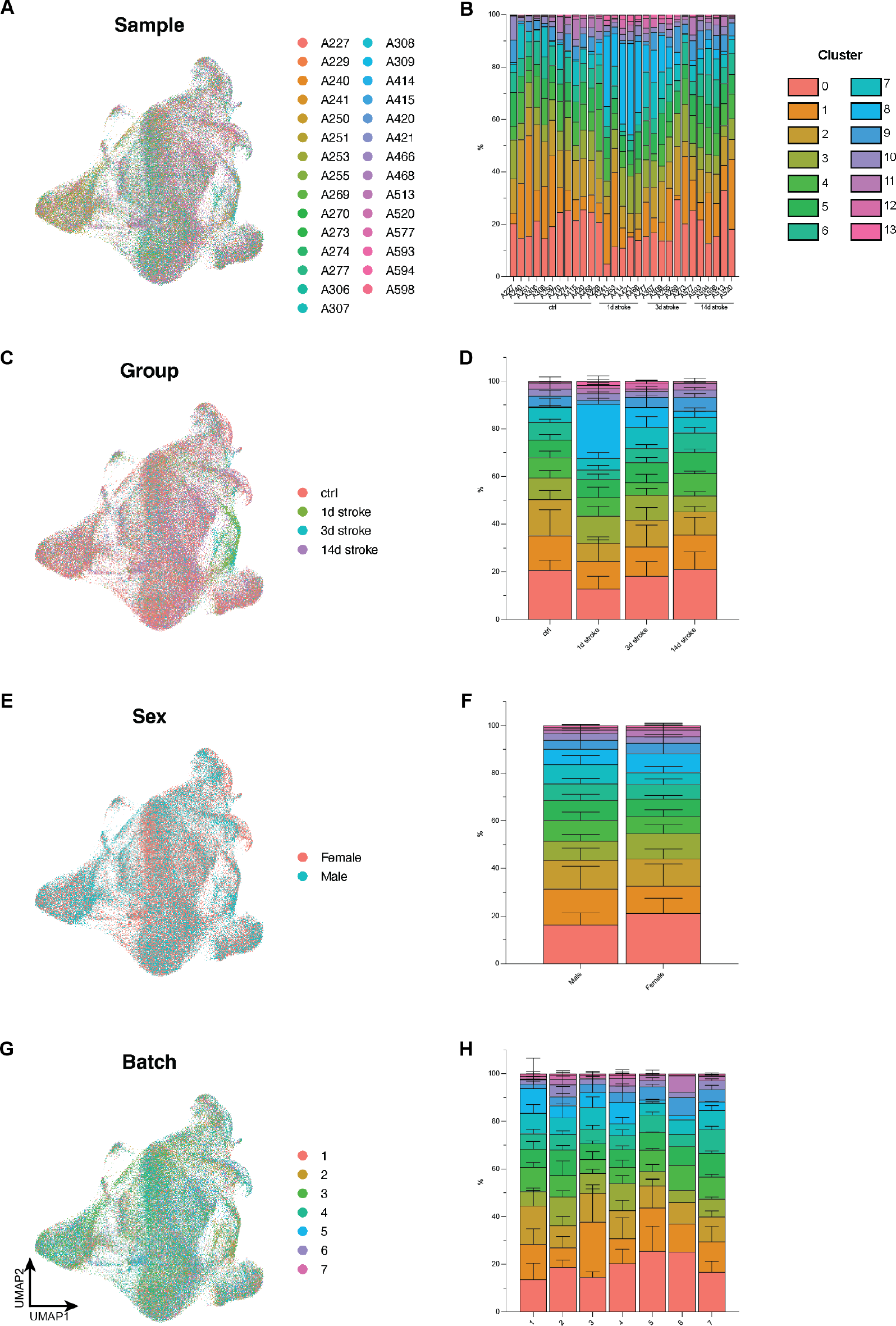
No significant differences in cluster composition across individual samples, experimental group, sex, or experimental batch. **(A)** UMAP of all astrocytes, with colors depicting individual samples. **(B)** Stacked bar graph showing % cluster composition in individual animals. **(C)** UMAP of all astrocytes, with colors depicting experimental condition. **(D)** Stacked bar graph showing % cluster composition in each experimental condition. **(E)** UMAP of all astrocytes, with colors depicting sex. **(F)** Stacked bar graph showing % cluster composition in each sex. **(G)** UMAP of all astrocytes, with colors depicting experimental batch. **(H)** Stacked bar graph showing % cluster composition in experimental batch.

**Figure S4.**
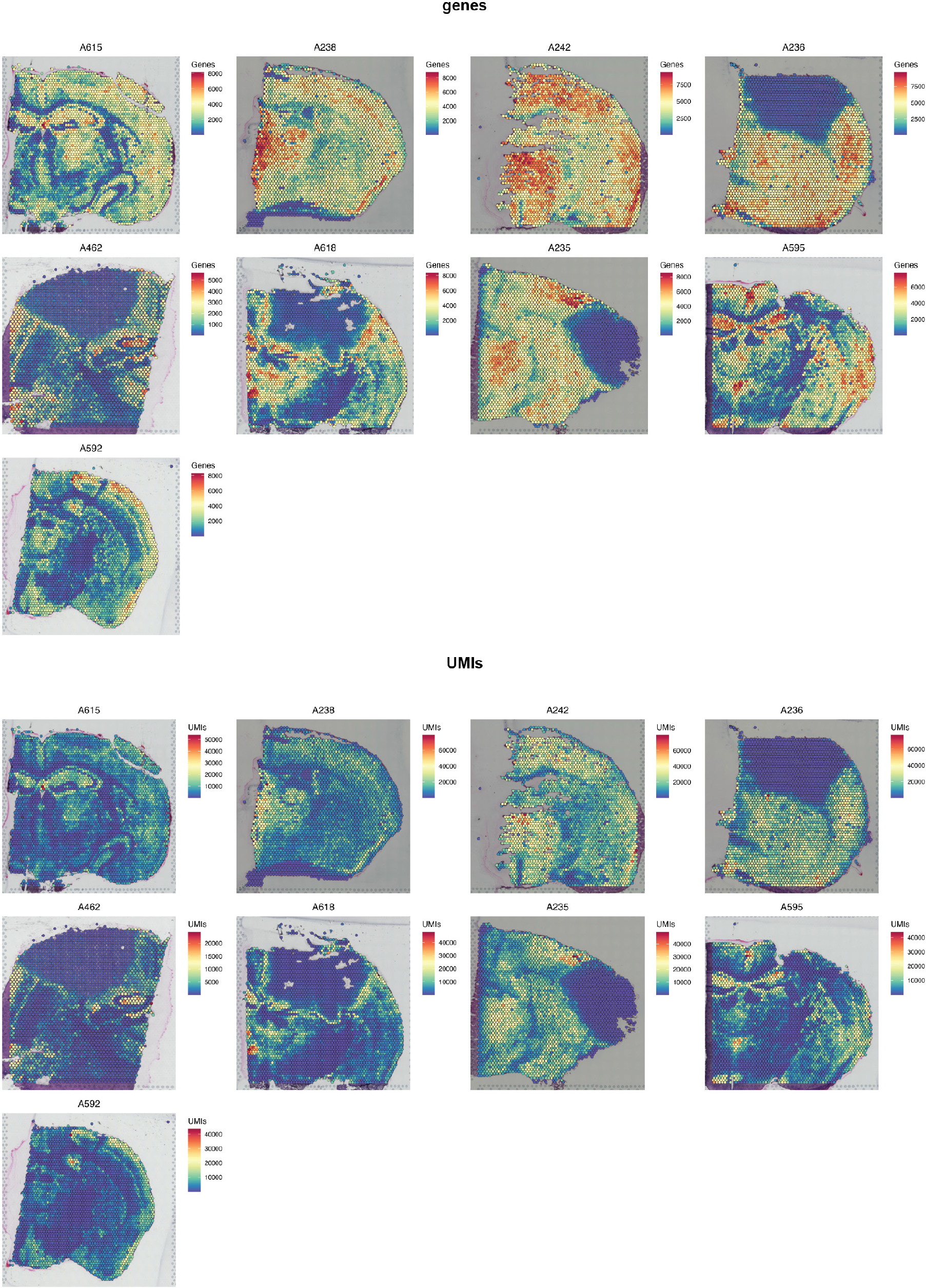
QC of spatial transcriptomics from control and 1, 3, and 14d post stroke tissue.

**Figure S5.**
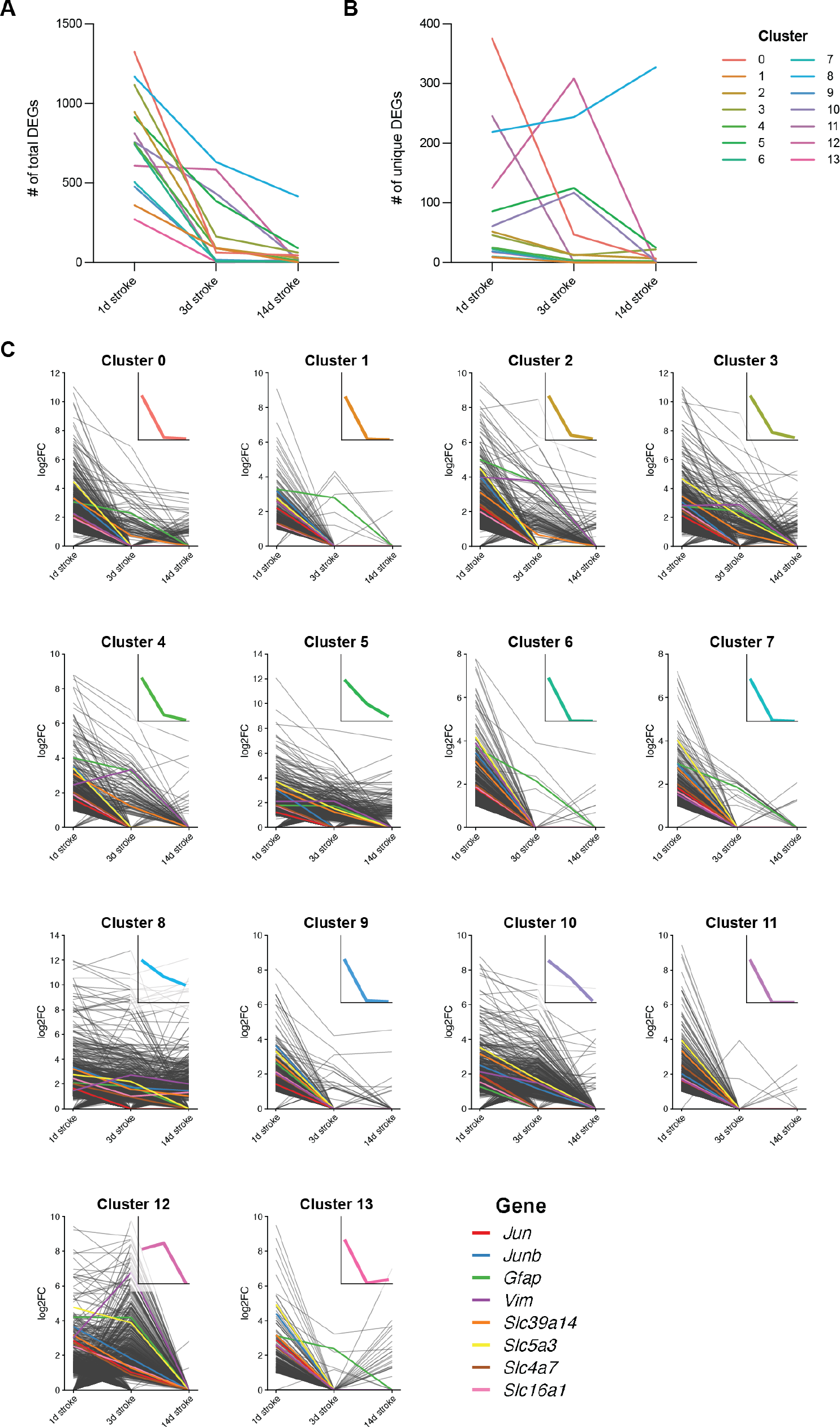
Dynamics of upregulated DEGs in each astrocyte cluster. **(A)** Number of total upregulated DEGs in each astrocyte cluster at each experimental timepoint. Lines colored by cluster. **(B)** Number of total upregulated DEGs in each astrocyte cluster at each experimental timepoint. Lines colored by cluster. **(C)** All upregulated DEGs by individual cluster at each timepoint, shown in gray. The individual genes discussed in Figure 4B (*Jun, Junb, Gfap, Vim, Slc39a4, Slc5a3, Slc4a7, Slc16a1*) are highlighted. Inset plots show the average fold change of upregulated genes in each cluster over time.

**Figure S6.**
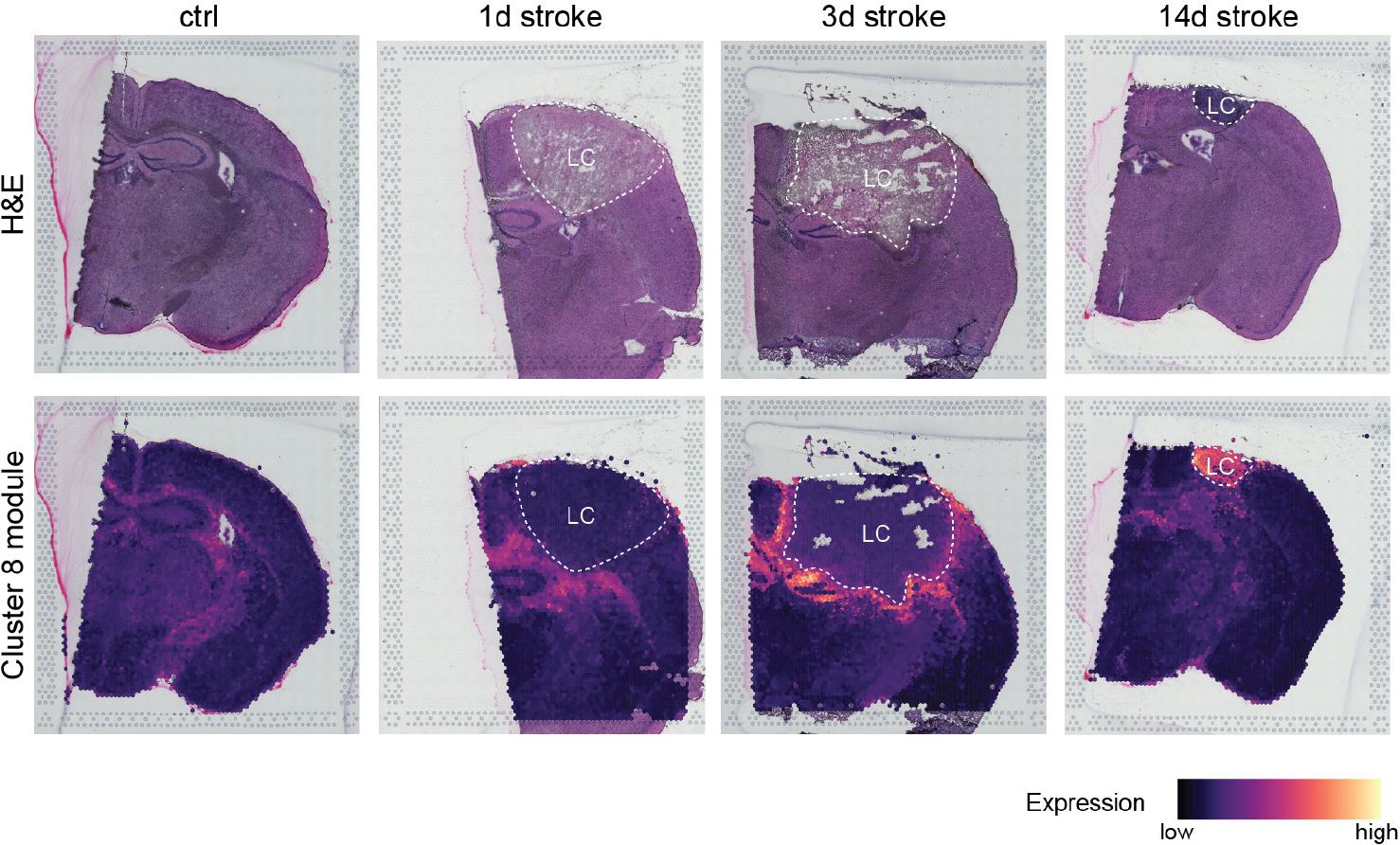
Representative Visium plots of cluster 8-enriched gene module for each experimental condition.

**Figure S7.**
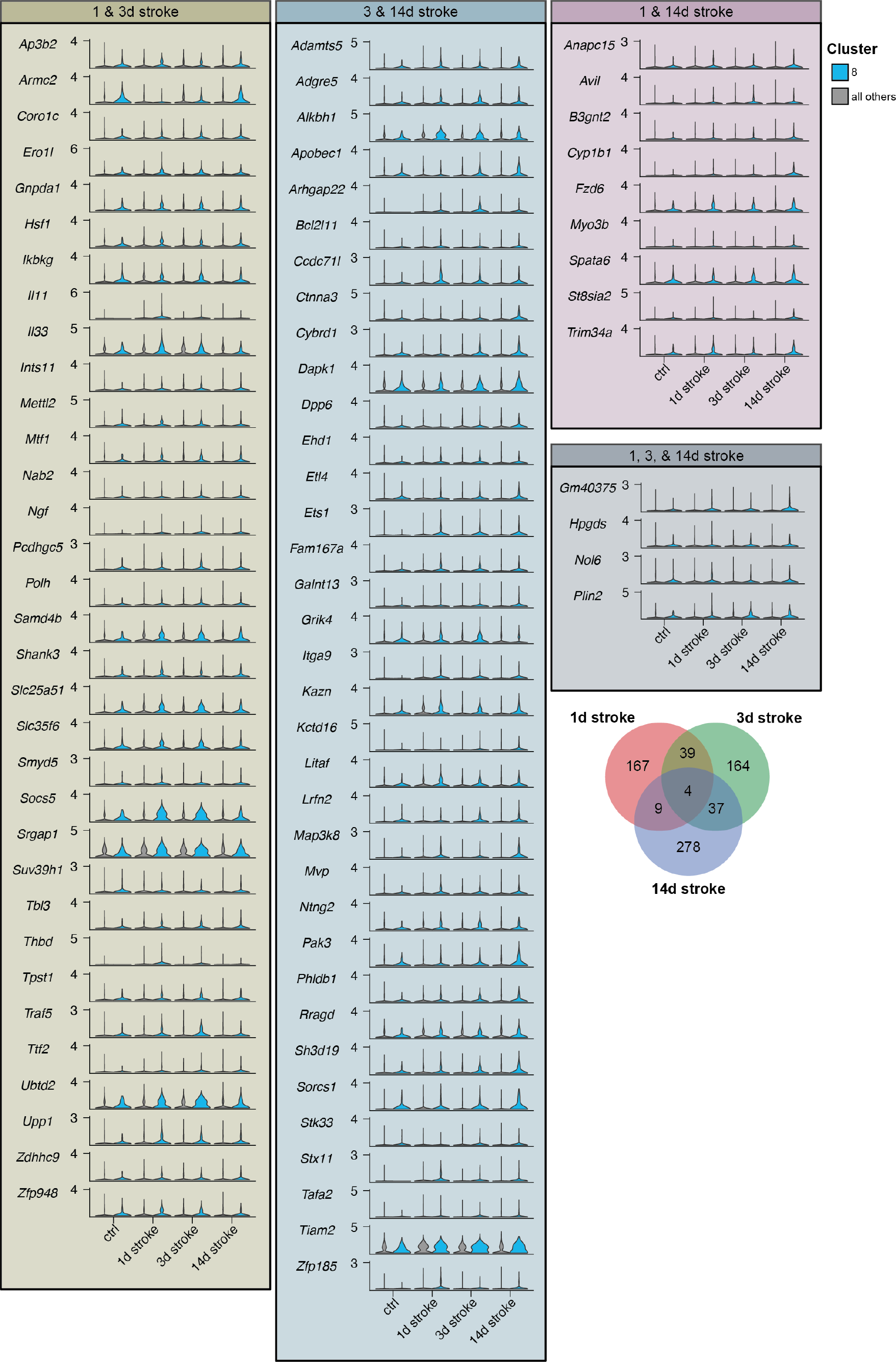
Violin plots of cluster 8 genes that are unique compared to all other clusters at their respective timepoints.

**Figure S8.**
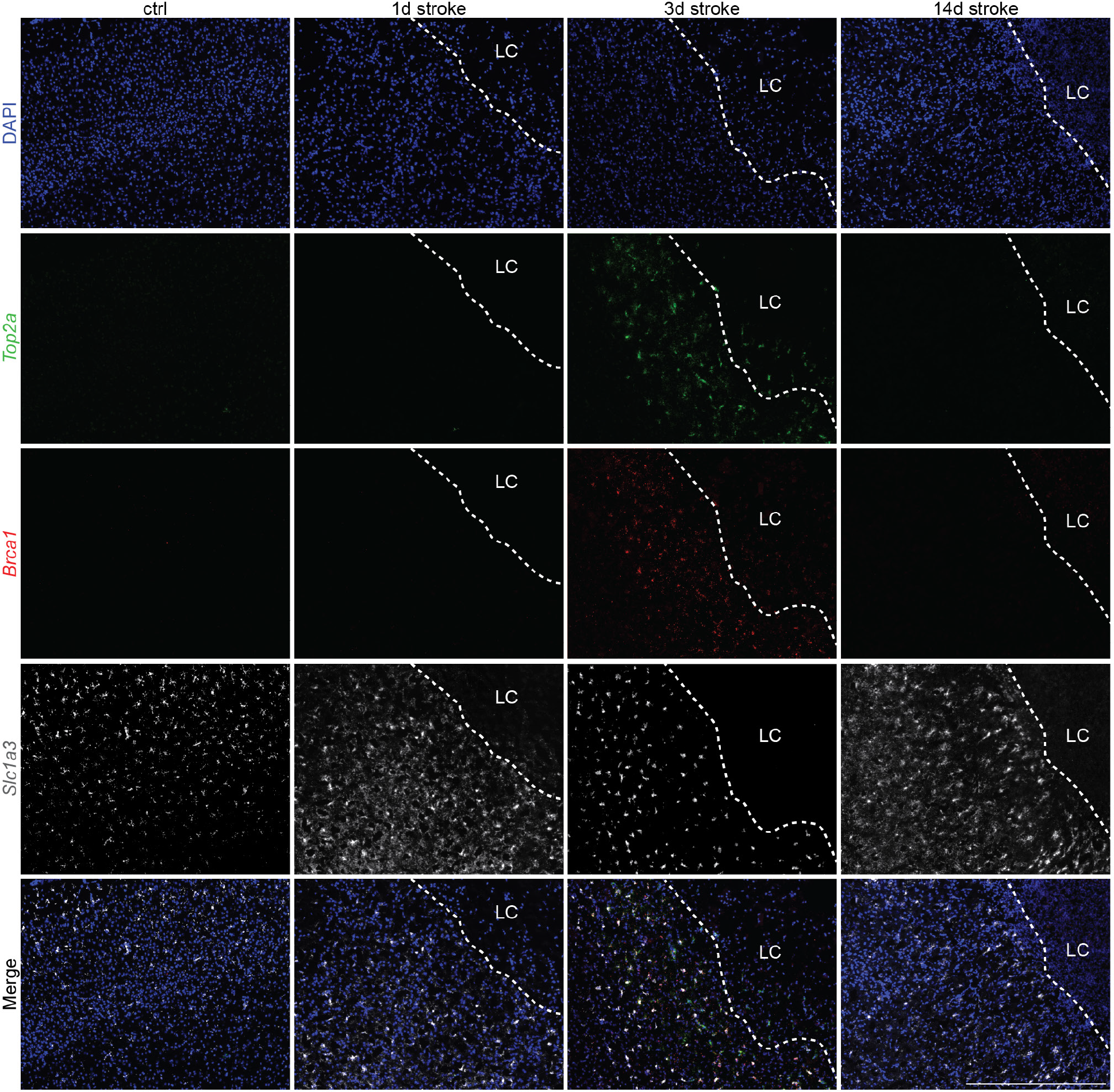
Expression of proliferative marker Top2a is restricted to 3d post stroke. Representative images of in situ hybridization for *Top2a, Brca1*, and *Slc1a3*, split by experimental condition. Scale = 500 μm; LC = lesion core.

**Figure S9.**
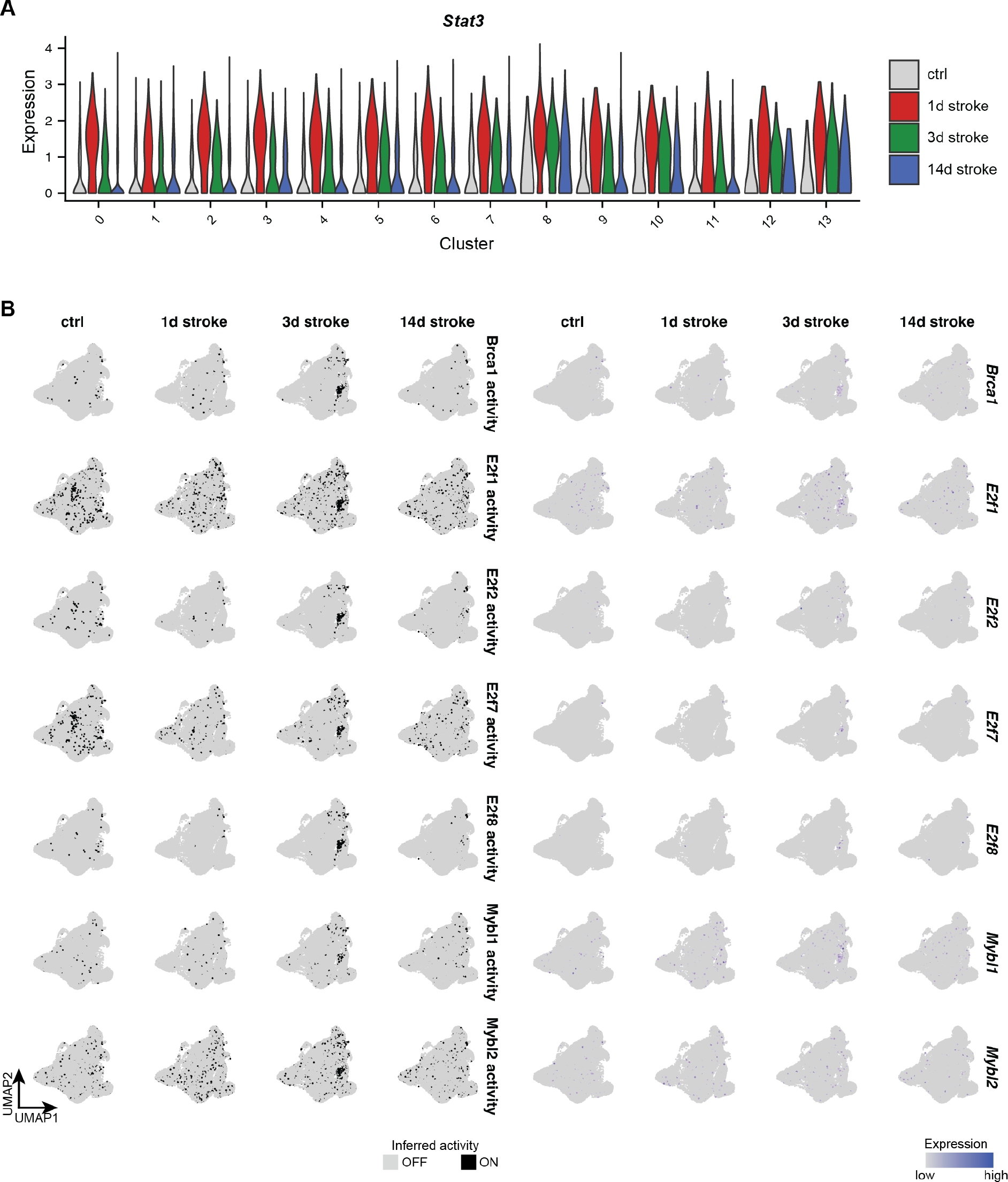
Potential transcriptional regulators of cluster 12 astrocytes. **(A)** Violin plots of Stat3 expression in each cluster, split by experimental condition. *Stat3* expression is upregulated not only in cluster 12 astrocytes following stroke, but also in many other astrocyte clusters. **(B)** Binarized plots for inferred transcriptional regulator activity with corresponding gene expression plots for potential cluster 12 transcriptional regulators identified with pySCENIC.

